# Rational engineering of a β-glucosidase (H0HC94) from glycosyl family I (GH1) to improve catalytic performance on cellobiose

**DOI:** 10.1101/2022.08.27.505235

**Authors:** Sauratej Sengupta, Pinaki Chanda, Bharat Manna, Supratim Datta

**Affiliations:** Protein Engineering Laboratory, Department of Biological Sciences, Indian Institute of Science Education and Research Kolkata, Mohanpur, West Bengal; Center for the Advanced Functional Materials, Indian Institute of Science Education and Research Kolkata, Mohanpur, West Bengal; Center for the Climate and Environmental Sciences, Indian Institute of Science Education and Research Kolkata, Mohanpur, West Bengal

**Keywords:** protein engineering, cellobiose, specific activity, enzyme specificity, molecular dynamics simulation, protein structure network

## Abstract

The conversion of lignocellulosic feedstocks by cellulases to glucose is a critical step in biofuel production. β-glucosidases catalyze the final step in cellulose breakdown, producing glucose, and is often the rate-limiting step in biomass hydrolysis. Rationally engineering previously characterized enzymes may be one strategy to increase catalytic activity and the efficiency of cellulose hydrolysis. The specific activity of most natural and engineered β-glucosidase is higher on the artificial substrate p-Nitrophenyl β-D-glucopyranoside (*p*NPGlc) than on the natural substrate, cellobiose. Based on our hypothesis of increasing catalytic activity by reducing the interaction of residues present near the active site tunnel entrance with glucose without disturbing any existing interactions with cellobiose, we report an engineered β-glucosidase (Q319A H0HC94) with a 1.8-fold specific activity increase (366.3 ± 36 µmol/min^/^mg), an almost 1.5-fold increase in *k*_*cat*_ (340.8 ± 27 s^-1^), and a 3-fold increase in Q319A H0HC94 cellobiose specificity (236.65 mM^-1^ s^-1^) over HOHC94. Molecular dynamic simulations and protein structure network analysis indicate that Q319A significantly increased the dynamically stable communities and hub residues, leading to a change in enzyme conformation and higher enzymatic activity. This study shows the impact of rational engineering of non-conserved residue to increase β-glucosidase substrate accessibility and enzyme specificity.

**TOC:** A rationally engineered β-glucosidase with a 1.5-fold increase in *k*_cat_, and a 3-fold increase in cellobiose specificity over the wild-type

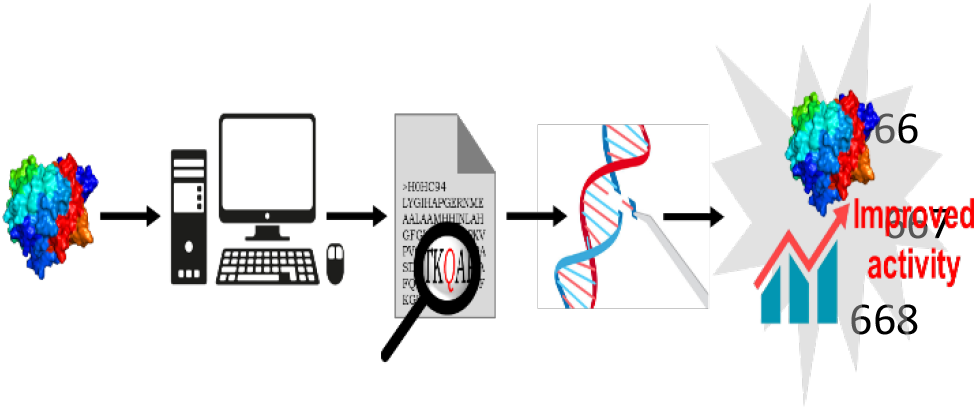

## Introduction

Production of biofuels from glucose produced through the saccharification of lignocellulosic biomass is a green alternative to fossil fuels pumped out of the earth. Cellulose is a major component of the naturally abundant lignocellulosic biomass and a homopolymer of glucose. Cellulases hydrolyze the cellulose to produce glucose. The cellulase cocktail is made up of at least three hydrolytic enzymes, namely, endoglucanase (EG), cellobiohydrolase (CBH), and β-glucosidase (BG). EG cleaves the β-1,4-glycosidic linkage of cellulose polymers to generate short-chain oligosaccharides, which CBH hydrolyzes to produce cellobiose units. Finally, BG’s cleave the β-1,4-glycosidic bonds to generate glucose that microbes can ferment to make biofuels.^1-3^ Most β-glucosidases that are reported in the literature are characterized by the model chromogenic substrate, *p*-Nitrophenyl beta-D-glucopyranoside (*p*NPGlc). Previously several attempts have been made to increase the activity of β-glucosidases.^4-7^ In most studies, the activity of engineered β-glucosidases increased more on the artificial substrate *p*NPGlc than the natural substrate, cellobiose.^4, 5, 8, 9^ The objectives of most such studies were to improve glucose tolerance,^6, 7, 10-12^, and thermal/pH stability.^4, 10^ Though a few studies reported an increase in cellobiose activity, there is a general lack of targeted engineering approach reports to improve cellobiose activity.^7, 9, 12^

Previously we reported a wild-type β-glucosidase (H0HC94) from *Agrobacterium tumefaciens* strain 5A belonging to the glycoside hydrolase family I, with high enzymatic activity on the chromogenic β-glycosidic substrate *p*NPGlc, and moderate activity on its natural substrate, cellobiose (Clb).^8, 13^ Furthermore, the enzyme is stabilized by increasing glucose concentrations resulting in improved half-life and more tolerance to [C2mim]-based ionic liquids.^13^ These promising properties motivated us to probe the enzyme further to enhance cellobiose activity and specificity.

In this study, we hypothesized that reducing the binding interaction of glucose molecules to the regions in or around the active site tunnel might enable greater accessibility of cellobiose, leading to higher cellobiose activity. We have used an *in-silico* rational approach to determine the target for mutation. Choosing the residue to mutate for engineering a better β-glucosidase is challenging because both substrate and inhibitor interact with a few common residues and modulate substrate binding and catalytic activity. Autodock Vina was used to dock cellobiose (substrate) and glucose (inhibitor) separately to the active site tunnel region.^14^ We identified the non-conserved Q319 at the junction of a β-sheet and a loop near the active site tunnel entrance and mutated it to an Alanine. Here we report the results of our characterization of the Q319A mutant, particularly its enhanced cellobiose activity and specificity, along with molecular dynamics simulations and protein structure network analysis to understand the role of Q319A on cellobiose activity.

## Materials and Methods

### Chemicals

Bacterial host strains and plasmids were purchased from Merck Millipore (Billerica, USA). Reagent-grade chemicals were used in this study. Primers were synthesized by Eurofins (Bangalore, India). Chromatography columns were acquired from GE Healthcare, Marlborough, USA. 30 kDa cut-off Amicon-Ultra-15 protein concentrators were bought from EMD Millipore (Billerica, USA). The culture media and cellobiose were purchased from Sigma-Aldrich (St. Louis, USA). *p*NPGlc was acquired from TCI Chemicals (Fukaya, Japan). All plastic consumables were purchased from Tarsons (Kolkata, India).

### Bacterial host, vectors, and media

pET-21b(+) (Novagen, Madison, USA) vector was used for cloning and expression of the mutant. *Escherichia coli* Top10F strain cells were used as the cloning host and *E. coli* BL21(DE3) (Stratagene Cloning Systems, La Jolla, CA) as the expression host. T7 RNA polymerase promoter was used for the overexpression of mutant protein by IPTG (G-Biosciences, St. Louis, USA) induction. All cells were screened and grown in Luria-Bertani agar/broth media bought from Sigma-Aldrich (St. Louis, USA).

### Molecular docking of cellobiose to H0HC94

Autodock Vina was used for the molecular docking of cellobiose and glucose to the active site tunnel of the protein (PDB: 6RJO).^14, 15^ In the configuration file, the dimension of the grid box used for docking was 40×40×40 covering the active site tunnel with a spacing of 1 Ǻ. Energy range and exhaustiveness parameter values were 4 and 8, respectively. Discovery studio generated 2D interaction diagrams of the wild-type protein with ligands such as cellobiose and glucose.^16^ A detailed 2D interaction diagram was generated by the PoseView tool in ProteinsPlus Server (proteins.plus/).^17^ An in-house python script was used to analyze the different poses generated by molecular docking and identify residues that most frequently interacted with glucose molecules. A histogram showing how often a residue interacted with glucose in various poses was generated with the data.

### Primer design, and PCR

The Q319A mutant was generated using the megaprimer-based PCR mutagenesis,^18^ using vector-specific T7 forward and T7 reverse primers along with the mutagenic primer. The mutagenic primer sequence containing flanking restriction sites, *Xho*I and *Hind*III, is 5’CCCTGCCACCAAAGCGGCCCCGGCCGTCAGC 3’. A 1 % Agarose gel was used to resolve and visualize the amplified gene product, and the amplified products were by QIAquick gel extraction kit (Qiagen, New Delhi, India).

### Cloning

The purified mutant gene was digested with *Xho*I and *Hind*III in CutSmart buffer (New England Biolabs, Ipswich, MA, USA). The digested gene product was ligated to the pre-digested and phosphatase-treated pET21b (+) plasmid by T4 ligase (New England Biolabs, Ipswich, MA, USA). Top10F *Escherichia coli* cells were transformed with ligated product using the heat shock method and used as the cloning host. The mutation was confirmed by Sanger sequencing at the DBS, IISER Kolkata sequencing facility.

### Expression and Protein Purification

Q319A H0HC94 was expressed and purified as described in our earlier study.^13^ The protein purity was confirmed by 10 % SDS–PAGE.

### Differential Scanning Fluorimetry (DSF)

Q319A H0HC94 DSF was performed, and the data were analyzed according to the previously reported protocol.^13^

### Circular Dichroism (CD)

The spectra were measured on a JASCO J-1500 Circular Dichroism Spectrophotometer (Easton, MA, USA) using a 1 mm cuvette. The scanning speed and bandwidth were 100 nm/min and 1 nm, respectively. The CD analysis was performed at room temperature (25 °C) with 1.9 µM wild-type and Q319A mutant in 50 mM potassium phosphate buffer, pH 7.4.

### Biochemical Characterization and Half-life Determination

The optimum temperature (T_opt_) and optimum pH (pH_opt_) of the mutant were ascertained by comparing the relative activity of the mutant between pH 4-10 (at 49 °C) and at 37 to 55 °C (at pH 7) respectively. 100 μL reactions were performed in 50 mM potassium phosphate buffer, using 0.05 μg enzyme, 20 mM *p*-nitrophenyl beta-D-glucopyranoside (*p*NPGlc) as the substrate, for a total reaction time of 30 min. The temperature and pH at which the mutant had the highest relative specific activity were chosen as the optimum reaction parameters for further kinetic analysis.

For half-life measurement, the protein was incubated in 50 mM potassium phosphate buffer, pH 7.0 at 49 °C up to 90 min. At regular intervals, reaction aliquots were taken out to determine enzyme specific activity. The relative specific activity was used to ascertain the half-life of the mutant protein using GraphPad Prism 9 software (GraphPad Software, La Jolla, CA) by fitting the data to a one-phase decay function. All experiments were performed in triplicates and repeated at least twice. The standard deviations among the repeats were below 10 %.

### Kinetic Parameters

Michaelis-Menten kinetics were performed on *p*NPGlc and cellobiose, with 0.5 mM to 40 mM substrate in 50 mM potassium phosphate buffer, pH 7, at 49 °C. In a 100 μL reaction volume, 0.16 μg of protein was used, and the reaction was quenched after 5 min. The buffer and the substrate were pre-incubated at 49 °C for 5 min before each reaction. For the *p*NPGlc assay, the reaction was stopped using 100 μL 0.4 M glycine (pH 10.8), and the reaction mixture was 20-fold diluted before taking an absorbance measurement at 405 nm in a clear bottom 96-well plate (Tarsons, Kolkata, India). The absorbance was measured by a Spectramax M2 spectrophotometer (Molecular Devices, San Jose, CA, USA). The experiment was done in triplicates and repeated thrice. The standard deviations among the repeats were below 10 %.

For the cellobiose reaction, the reaction was stopped by heat inactivation at 95 °C for 15 min. The reaction product was 8 -fold diluted in 50 mM potassium phosphate buffer, pH 7.0, before performing the GOD-POD assay (Sigma, St Louis, USA). The absorbance of the final product mixture was recorded at 527 nm as described for the reaction on *p*NPGlc. GraphPad Prism 9 software (GraphPad Software, La Jolla, CA) was used to analyze the velocity data and calculate the kinetic constants such as *K*_*m*_ and *k*_*cat*_. The experiment was done in duplicate and repeated twice with different batches of purified protein. The standard deviations among the repeats were below 10 %.

### Molecular dynamics (MD) simulation

MD simulations were performed using the X-ray crystal structure (PDB ID: 6RJO) of β-glucosidase from *A. tumefaciens* 5A.^15^ AMBER20^19^ was utilized to carry out the simulations. The residue numbering of the enzyme is as per previous reports^20, 21^. The AMBERff14SB^22^ force field was employed for the protein, whereas the GLYCAM06^23^ and General Amber Force Field (GAFF) were assigned to cellobiose and *p*NPGlc molecules, respectively. The Q319A mutant was prepared from the WT by using Chimera.^24^ Four systems were prepared for the MD simulations: S1: WT+0.02 M *p*NPGlc, S2: WT+0.02 M Cellobiose, S3: Q319A+0.02 M *p*NPGlc, S4: Q319A+0.02 M Cellobiose (Supplementary Table S1). The simulation systems were built using PACKMOL.^25^ The enzymes were protonated at their respective pH_opt_ using the PDB2PQR^26^ server (Supplementary Table S1). The TIP3P water model’s parameters were adopted for the water molecules^27^. The bulk system’s characteristics were simulated using periodic boundary conditions (PBC). The particle mesh Ewald (PME) summation method was used to calculate the electrostatic interaction with the distance threshold for nonbonded interactions of 10 Å.^28^ With the Langevin thermostat, the temperature was kept at the corresponding T_opt_ of the enzyme (Supplementary Table S1). The SHAKE method was used to constrain Hydrogen atom containing bonds. Using the steepest descent algorithm, the starting geometries of all systems were energy minimized for 10,000 steps. With a time-step of 2 femtosecond (fs), the MD production run was carried out for 200 ns under the NPT ensemble. The CPPTRAJ^29^ module was used to evaluate the trajectory data, which was saved with a 1 picosecond (ps) interval. The simulation results were displayed using Visual molecular dynamics (VMD).^30^

### Protein Structural Network (PSN) analysis

A dynamic protein structure network (PSN) was constructed to examine protein structures generated from simulation trajectories. The protein structures from each frame of the simulation were utilized to build the PSN, where the amino acid residues served as nodes, and edges were formed between two nodes involved in non-covalent interactions. To determine the strength of the edges, the percentage interaction (I_ij_) between residues i and j is:

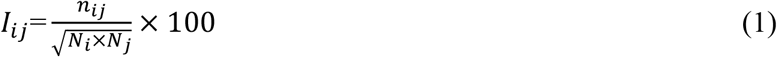

where n_ij_ is the number of atom pairs between residues i and j within 4.5 Å.^20^ The normalization factors N_i_ and N_j_ are unique to the types of residues i and j, respectively. A community is a union of cliques (complete connected subgraph) that share common nodes.^20, 31^ The nodes with at least four edges were attributed as hub residues.^32^ All the PSN analyses were performed using PSN Tools.^33^

## Results and Discussion

### Selection of mutation site (Q319)

Molecular docking results indicate that residues such as Q25, H126, N170, Y299, W405, and E359 (catalytic residue), interact with cellobiose (Supplementary Figure S1a). Similar interactions were also reported in the crystal structure (PDB ID: 6RJO), where the ligand was salicin (a substrate analog).^15^ Mutating such residues might disrupt the substrate-enzyme interactions and affect biocatalysis. Therefore our goal was to introduce a mutation that would disrupt interactions between active site tunnel residues and glucose without affecting any interactions with cellobiose. Glucose interacts with the backbone of amino acid residues (instead of the side chain) such as T300, G331, and P301, so mutating these residues might not affect the glucose binding. Other residues like W127, W173, C174, L178, H185, H229, S230, N227, N297, Y298, M302, R303, Q319, A322, K327, W332, E333, E359, E412, W413, F421 interacts with glucose through its side-chain by forming hydrogen bonds and/or van der Waal interactions (Supplementary Figure S1b). We preferred to disrupt the stronger bonding interactions over van der Waal interactions. Q25, E171, C174, N227, H229, T300, P301, R303, Q319, K327, G331, E333, E359, W405, E412, and W41 formed hydrogen bonds with glucose. Of these, Q25, E171, T300, Q319, K327, and W405 appeared at least twice while searching through the different poses of glucose binding generated by molecular docking (Supplementary Figure S1). Q319 appeared in two poses to form hydrogen bonds with glucose, does not interact with cellobiose (Supplementary Figure S2), and is a non-conserved residue, so we selected Q319 to study its effect on glucose binding and catalysis.

### Engineering of Q319A H0HC94

We performed site-directed mutagenesis on the wild-type gene to produce the Q319A HOHC94. To compare its properties with the wild-type (WT), we overexpressed the enzyme in *E. coli* and purified it (Supplementary Figure S3a, S3b). SDS PAGE of the purified mutant protein showed a clear, prominent band at approximately 52 kDa (Supplementary Figure S3c). The optimum temperature (T_opt_) and optimum pH (pH_opt_) of the Q319A mutant were 49 °C (Figure 1a) and 7.0 (Figure 1b), respectively, compared to 52 °C and 7.2, respectively, of the WT.^13^ Thus, while the pH_opt_ was almost unchanged, the T_opt_ decreased by 3 °C.

**Figure 1.**
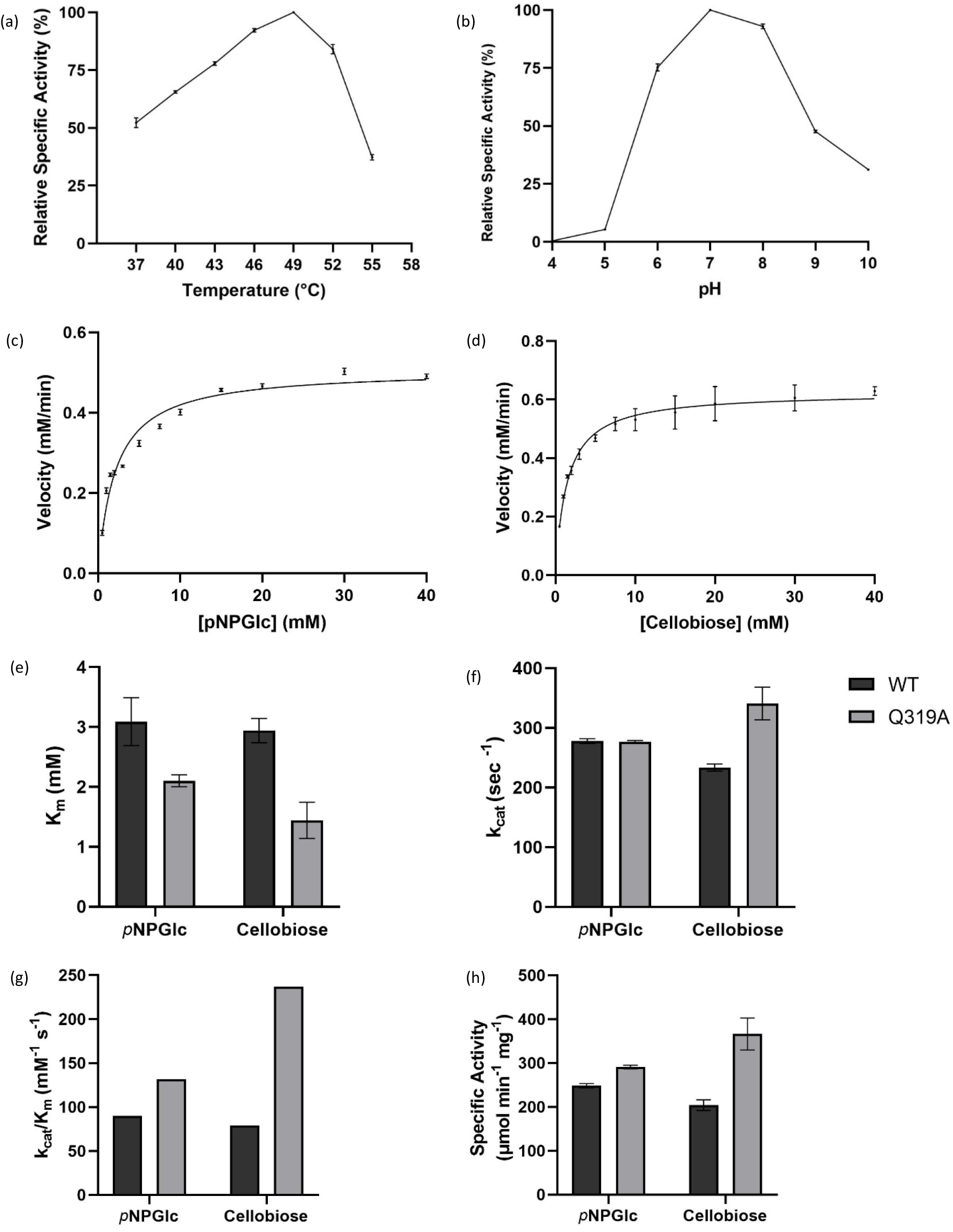
**(a)** Relative activity profile of Q319A H0HC94 mutant between 37°C to 55° C in 50 mM potassium phosphate buffer pH 7; **(b)** Relative activity profile of Q319A H0HC94 mutant through pH 4-10 in 50 mM potassium phosphate buffer at 49 °C; **(c)** Michaelis-Menten kinetic profile of Q319A H0HC94 mutant on *p*-nitrophenyl beta-D-glucopyranoside (*p*NPGlc) in the absence of glucose using 0.16 μg of enzyme in 50 mM potassium phosphate buffer, pH 7, at 49 °C; **(d)** Michaelis-Menten kinetic profile of Q319A H0HC94 mutant on cellobiose, in the absence of glucose using 0.16 μg of enzyme in 50 mM potassium phosphate buffer, pH 7, at 49 °C; **(e)** Comparison of K_m_ between WT and Q319A on *p*NPGlc; **(f)** Comparison of *k*_*cat*_ between WT and Q319A on *p*NPGlc; (g) Comparison of *k*_cat_*/K*_M_ between WT and Q319A on *p*NPGlc; **(h)** Comparison of specific activity of WT and Q319A on *p*NPGlc and cellobiose.

### Thermal stability of the mutant

An enzyme’s structural and thermal stability often plays a crucial role in its catalytic activity. Differential Scanning Fluorimetry (DSF) analysis was used to measure melting temperature (T_m_) to understand the effect of mutation on enzyme stability. The T_m_ of the Q319A mutant was 53.9 °C (Figure 2c), similar to H0HC94 T_m_ of 53.7 °C.^13^ To verify the absence of any overall structural distortion due to the mutation, we measured the circular dichroism (CD) spectra at room temperature and observed no significant difference between the secondary structure of wild-type H0HC94 and Q319A (Figure 2b, Supplementary Table S2).

**Figure 2.**
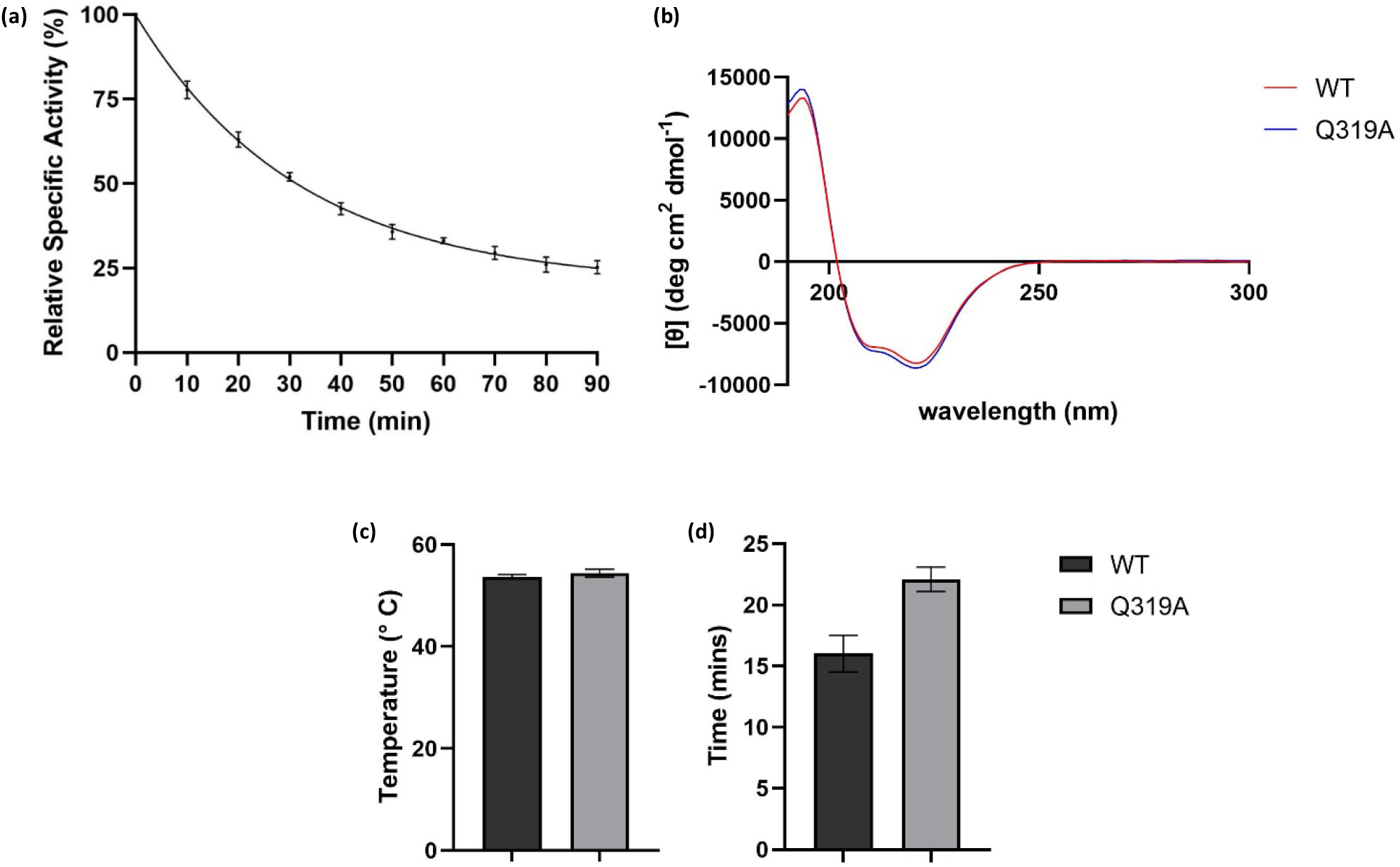
**(a)** Half-life of Q319A H0HC94. The enzyme in 50 mM potassium buffer, pH 7, was incubated at 49 °C, and aliquots were removed at 10 minute intervals for specific activity measurement. **(b)** Circular Dichroism spectra of WT H0HC94 and Q319A (1.9 μM) in 50 mM potassium phosphate buffer, pH 7 at room temperature (25 °C) **(c)** Comparison of melting temperature of WT and Q319A determined by differential scanning fluorimetry (DSF) and CD **(d)** Comparison of the half-life of WT and Q319A at their optimum temperatures.

The half-life of Q319A H0HC94 increased by 1.4-fold compared to the wild-type. The half-life of Q319A was 22 ± 1 min (Figure 2a) compared to 16 min of the wild-type^13^. This increase suggests a slight enhancement in Q319A thermal stability.

### The catalytic efficiency of Q319A compared to the WT

Specific activity on *p*NPGlc increased 1.2-fold from 248 µmol/min/mg of the WT to 291 ± 4 µmol^-^min^-1^ mg^-1^ for the Q319A mutant (Figure 1h). The analysis of kinetic velocity data using *p*NPGlc as substrate (Figure 1c) shows that Q319A has a lower *K*_m_ of 2.1 ± 0.1 mM compared to 3.09 ± 0.4 mM of the WT (Figure 1e).^13^ The lower *K*_m_ probably reflects an increased affinity of the substrate due to its enhanced accessibility to the substrate binding site. The turnover number, *k*_cat_, on *p*NPGlc, at 276.6 ± 2.3 s^-1^, almost remained the same as the wild-type (277.9 ± 4 s^-1^, Figure 1f).

Michaelis Menten kinetic assays of Q319A H0HC94 were also performed on its natural substrate, cellobiose (Figure 1d). The specific activity on cellobiose increased 1.8-fold from 204 ± 12 µmol/min/mg to 366.3 ± 36 µmol/min/mg (Figure 1h). The *K*_m_ decreased from 2.94 mM to 1.44 ± 0.3 mM (Figure 1e), while the *k*_cat_ increased approximately 1.5-fold from 233.4 ± 6 s^-1^ to 340.8 ± 27 s^-1^ (Figure 1f). Thus, the substrate specificity of Q319A increased 3-fold to 236.65 mM^-1^ s^-1^ from 79.32 mM^-1^ s^-1^ of the WT (Figure 1g). To make sense of the increase, we compared previous reports of engineered GH1 β-glucosidase in the literature (Table 1). A β-glucosidase, BglA, from *Caldicellulosiruptor saccharolyticus* (CsBglA) was engineered to improve catalysis at a lower temperature.^9^ By random mutagenesis, the *k*_cat_/*K*_m_ of a triple mutant variant of a glucose tolerant Bgl6 was improved 3-fold, from 0.56 mM^-1^ min^-1^ to 1.69 mM^-1^ min^-1^.^11^ Recently, in another study, the authors engineered a glucose tolerant variant of Bgl15 and enhanced cellobiose specificity from 0.10 ± 0.01 to 1.20 ± 0.09 mM^-1^sec^-1^.^12^ While the cellobiose specificity reported in this work is higher than previously reported, current efforts are on to further improve substrate specificity similar to that of Ks5A7 (Table 1).

**Table 1.** A comparison of Q319A optimum temperature, specific activity, *K*_m_, *k*_cat_, and *k*_cat_/*K*_m_ on cellobiose with WT and engineered and naturally available GH1 β-glucosidases previously reported in the literature.

| Enzyme | Optimum temperature (°C) | Specific activity ($\mu\text{mol min}^{-1} \text{mg}^{-1}$) | $K_m$ (mM) | $k_{cat}$ ($\text{s}^{-1}$) | $k_{cat}/K_m$ ($\text{mM}^{-1} \text{s}^{-1}$) | Reference |
| --- | --- | --- | --- | --- | --- | --- |
| H0HC94 | 52 | $204 \pm 12$ | 2.94 | $233.4 \pm 6$ | 79.32 | 8 |
| Q319A H0HC94 | 49 | $366.3 \pm 36.6$ | $1.44 \pm 0.3$ | $340.8 \pm 27.4$ | 236.65 | This study |
| CsBglA | 80 | $359 \pm 10$ | $61.15 \pm 10.53$ | $328 \pm 26.2$ | $5.4 \pm 1.4$ | 9 |
| CsBglA-LYTH | 65 | $552 \pm 40$ | $37.06 \pm 6.87$ | $504 \pm 36.7$ | $13.6 \pm 3.5$ | 9 |
| Bgl 1317 | 60 | $2.16 \pm 0.07$ | ND | ND | ND | 7 |
| A397R Bgl 1317 | ND | 3.7* | ND | ND | ND | 7 |
| Bgl6 | 50-55 | $21.71 \pm 0.27$ | $38.45 \pm 1.51$ | $21.50 \pm 0.38$ | 0.56 | 11 |
| Bgl6 M3 | 60 | ND | $49.19 \pm 1.75$ | $83.11 \pm 4.12$ | 1.69 | 11 |
| Bgl15 | 50 | ND | $2901.81 \pm 72.50$ | $292.82 \pm 16.11$ | $0.10 \pm 0.01$ | 12 |
| Bgl15 5R1 | 60 | ND | $57.26 \pm 4.32$ | $68.51 \pm 2.75$ | $1.20 \pm 0.09$ | 12 |
| Cel3B | 60 | $33 \pm 3.6$ | 0.31 | $6.7 \pm 0.83$ | 21.6 | 34 |
| HiBgl3C | 60 | 56 | 6.63 | 152.5* | 23 | 35 |
| HGT-BT | 50 | 353 | 7 | 253* | 36 | 36 |
| Recombinant Ks5A7 from Kusaya gravy metagenome | 50 | $170 \pm 20$ | $0.358 \pm 0.055$ | 138 | 386 | 37 |

\*Calculated using the parameters reported in the papers

### Molecular dynamic insights into the structural stability of Q319A H0HC94

H0HC94 contains (β/α)_8_-barrel structural fold and belongs to the GH1 family (Supplementary Figure S4).^15^ The dynamic properties of WT H0HC94 and the Q319A variant were assessed by performing all-atom molecular dynamics simulations in the presence of the chromogenic substrate *p*NPGlc and natural substrate, cellobiose (Supplementary Table S1). Root-Mean-Square Deviation (rmsd) was calculated based on the backbone C_α_ atoms. The initial structure was considered as a reference. The average rmsd values were 1.54 ± 0.22 Å, 1.24 ± 0.13 Å, 1.17 ± 0.11 Å, 1.18 ± 0.11 Å in S1 (WT + 0.02 M *p*NPGlc), S2 (WT + 0.02 M Cellobiose), S3 (Q319A +0.02 M *p*NPGlc) and S4 (Q319A +0.02 M Cellobiose), respectively (Figure 3a). Low average rmsd values indicated that neither WT nor Q319A deviated from their starting structures, and the Q319A mutation did not cause any significant alteration to the H0HC94 structure at the backbone level during the simulation timescale. A similar rmsd pattern was previously reported, signifying the overall structural stability of the enzyme under the studied timescales.^20, 21^

**Figure 3.**
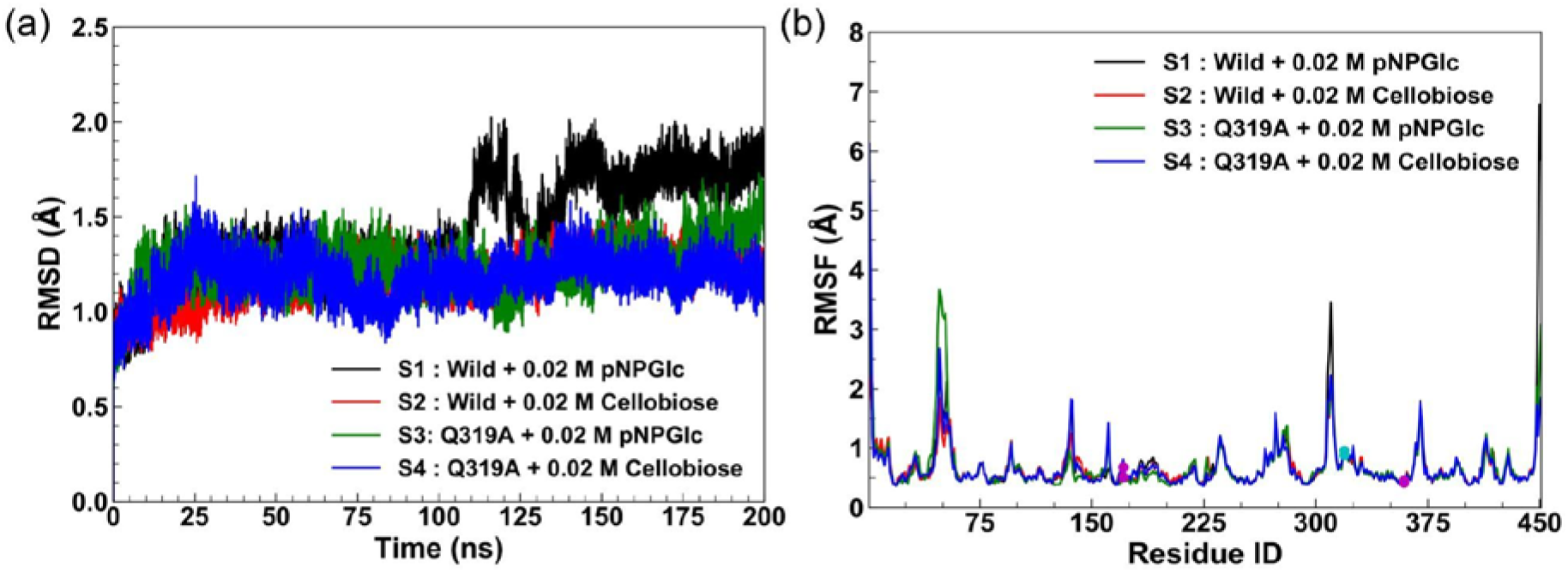
The structural changes of H0HC94 during MD simulations showing **(a)** Root-mean-square deviation (rmsd) of H0HC94 in S1 (WT+*p*NPGlc), S2 (WT+Cellobiose), S3 (Q319A+*p*NPGlc) and S4 (Q319A+Cellobiose) systems. **(b)** Root-mean-square fluctuations (rmsf) of the H0HC94 in S1 (WT+*p*NPGlc), S2 (WT+Cellobiose), S3 (Q319A+*p*NPGlc) and S4 (Q319A+Cellobiose) systems. The catalytic residues, acid/base residue E171, and nucleophilic residue E359 are marked in magenta circles. A cyan circle marks the mutated residue Q319A.

To understand the role of amino acid residues, the residue flexibility was measured by the Root-Mean-Square fluctuation (rmsf) profiles during the simulations (Figure 3b). The active site catalytic residues, active site tunnel residues, and gatekeeper residues exhibited low average rmsf (≤ 1.0734 Å) across the H0HC94 variants indicating no significant changes due to the Q319 mutation (Supplementary Table S3). However, there were slight fluctuations between the WT and Q319A in the presence of substrates. For example, due to Q319A mutation, residues E171, H126, W127, N170, C174, W331, W405, W413, L178, and H185 had slightly reduced flexibility, whereas the flexibility of E359, N297, N227, H229, Y299, and T300 slightly increased in the presence of *p*NPGlc (S3) compared to WT (S1). However, there were more residues with higher flexibilities (E171, E359, W127, N170, C174, N297, W413, L178, H185, Y299, and T300) and few with reduced fluctuations (H126, W331, W405, N227, and H229) in the presence of cellobiose compared to WT (S3). This differential local residue flexibility within the active site micro-environment suggests an alteration in enzyme-substrate interaction between the chromogenic substrate (*p*NPGlc) and natural substrate (cellobiose).

### Interaction between enzyme and substrate by Hydrogen bond analysis

During the simulation process, the interactions of both substrates with WT and Q319A were investigated to look for non-bonding driving forces like enzyme-substrate inter-hydrogen bonds. We previously reported that glucose might form multiple hydrogen bonds to H0HC94 through hydroxyl groups.^20, 21^ Our current simulations show that the average number of inter-hydrogen bonds between H0HC94 and substrates were 10 ± 3, 20 ± 4, 9 ± 3, and 15 ± 3 in S1, S2, S3, and S4, respectively (Supplementary Table S1; Figure 4a). Cellobiose formed comparatively more hydrogen bonds with both WT and mutant than *p*NPGlc. Interestingly, there was a slight decrease in the average number of hydrogen bonds in Q319A (both in *p*NPGlc and cellobiose) compared to the WT. It suggests that due to Q319A mutation, the interaction between the enzyme and substrate is slightly altered compared to the WT. To pinpoint the exact residue-specific interactions that were affected due to the mutation, the hydrogen bond fraction for each residue was computed in every system. The hydrogen bond fraction refers to the total amount of time a specific hydrogen bond was present throughout the entire simulation trajectory. There were 9, 12, 31, and 15 residues with hydrogen bonding fractions greater than 0.1 in S1, S2, S3, and S4, respectively (Figure 4b). Though the hydrogen bonding fraction was low, there were notable differences across the systems. For example, four active site tunnel residues (Q25, N297, E359, and W413) had hydrogen bonding fractions greater than 0.1 in S1, whereas there were 10 residues (E171, H185, N227, Y299, T300, D329, E359, W405, E412, and E415) inside the active site tunnel in S3, while enzymes in both the systems were interacting with *p*NPGlc. It indicates that the Q319A mutation enabled a preferential interaction between the active site tunnel residues and *p*NPGlc leading to higher *p*NPGlc activity of the mutant. Besides, 9 residues (E171, N227, H251, Y299, R303, Q319, E359, W405, E415) around and outside the active site tunnel were interacting with cellobiose in S2. The number of interacting residues was similar (Q25, E171, Y182, Y299, T317, E359, E412) in S4. As all the enzymes were interacting with their corresponding substrates (*p*NPGlc or cellobiose), the hydrogen bonding fraction with either of the two catalytic residues (E171, E359) was high in both WT and Q319A. Thus, the hydrogen bonding analyses revealed that the mode of interaction with *p*NPGlc or cellobiose was different in both WT H0HC94 and Q319A. Also, the Q319A mutation enhanced the active site tunnel hydrogen-bonding interactions with *p*NPGlc, while on the other hand, the total number of surface interactions was reduced in the case of the cellobiose. In both scenarios, the Q319A exhibited higher activity than the WT enzyme, as confirmed by experiments.

**Figure 4.**
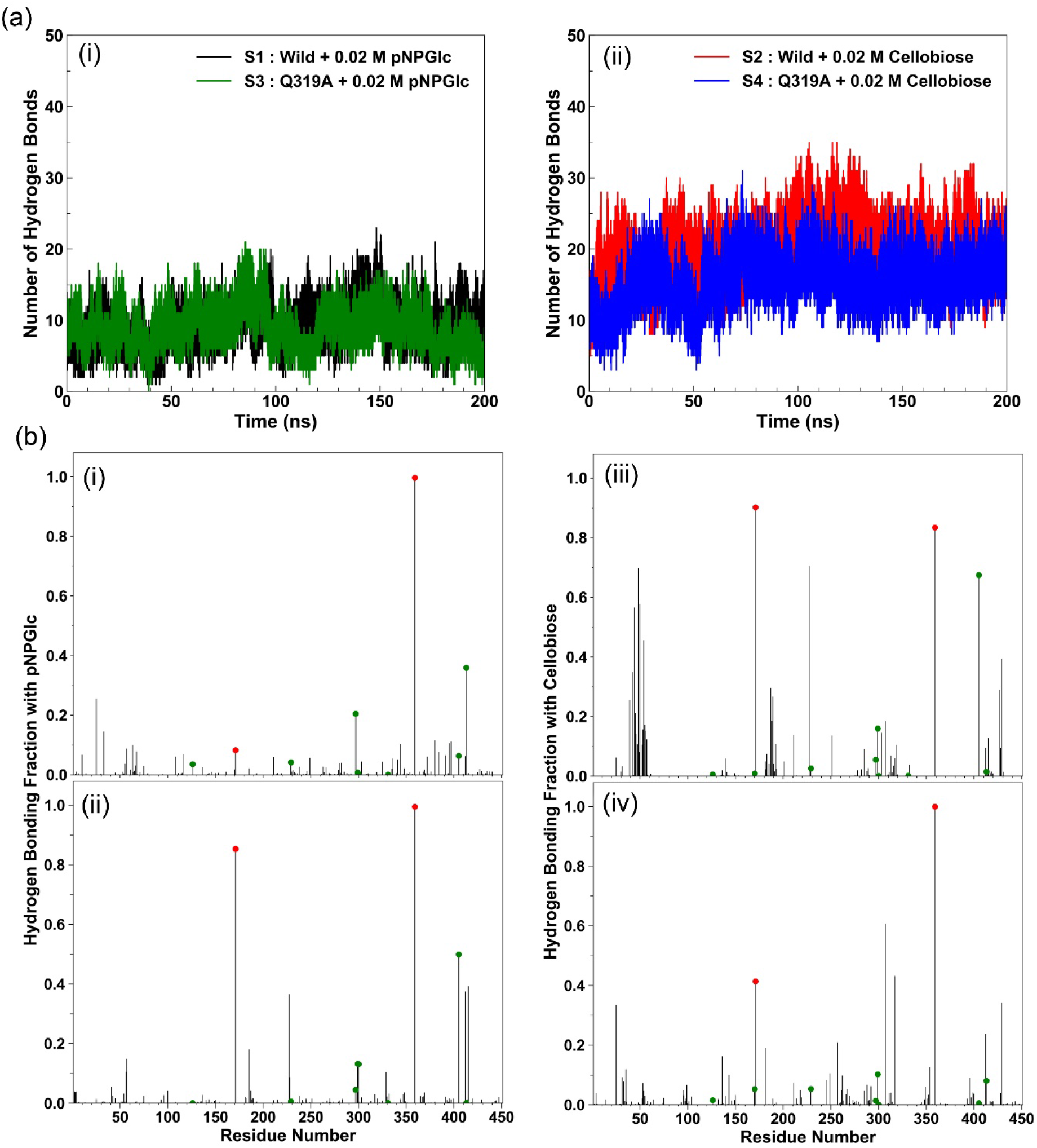
**(a)** The number of hydrogen bonds between H0HC94 and (i) pNPGlc and (ii) Cellobiose in S1 (WT+pNPGlc), S2 (WT+Cellobiose), S3 (Q319A+pNPGlc), and S4 (Q319A+Cellobiose) systems during the simulation; **(b)** The hydrogen bond fractions between H0HC94 and substrates (pNPGlc/ Cellobiose) in **(i)** S1 (WT+pNPGlc), **(ii)** S3 (Q319A+pNPGlc), **(iii)** S2 (WT+Cellobiose), and **(iv)** S4 (Q319A+Cellobiose) systems.

### Conformational change of the proteins using Protein Structure Network (PSN) Analysis

In recent years, dynamic changes in the conformation of enzymes from different glycoside hydrolases (GHs) such as endoglucanase Cel12A,^32^ xylanase,^31^ and β-Glucosidase^20, 21^ was investigated by PSN to provide global characteristics of protein structures during MD simulations. We examined the conformational changes of WT and Q319A in the presence of *p*NPGlc and cellobiose by constructing the PSN on each enzyme structure from the simulation trajectories and evaluating it using novel network parameters, *i*.*e*., dynamically stable network communities (connected cliques) and hubs (nodes with ≥ 4 edges). The communities/hubs are considered dynamically stable if they were present in more than 50 % of the simulation trajectories. The PSN analysis revealed considerable differences in the community (C) pattern of the H0HC94 in S1 (communities=6, nodes=43), S2 (communities=7, nodes=59), S3 (communities=5, nodes=67), and S4 (communities=6, nodes=61) systems (Figure 5, 6). Out of the 6 communities in S1 (WT+0.02 M *p*NPGlc), the active site community (C1) comprised of 20 residues (Figure 5a, 5b). On the other hand, the active site community (C1) in Q319A (S3), had increased to 54 residues (Figure 6a, 6b). Moreover, the total number of edges was increased, signifying the elevated interactions within the Q319A (S3) compared to the WT (S1). Also, the total number of hub residues increased in S3 (16) compared to S1 (12), indicating higher integrity of the PSN and increased stability of the mutant structure. The protein in S2 (WT+0.02 M Cellobiose) had the active site community (C1) comprised of 24 residues (Figure 5c, 5d). But, due to the Q319A mutation (S4), the active site community (C1) had increased to 45 residues (Figure 6c, 6d). Moreover, the total number of edges increased, signifying the elevated interactions within Q319A (S4) compared to WT (S2). Also, the total number of hub residues increased in S4 (15) compared to S2 (9), indicating a more compact PSN and stabilized Q319A structure. The PSN analysis reveals that the active site conformation was significantly remodeled due to the Q319A mutation. Substrate-induced conformational changes were different for *p*NPGlc and cellobiose. Thus, the PSN analysis reveals that both the active site structure and the conformation of Q319A are more stable and robust, corroborating the increase in experimentally determined activity and specificity over the WT.

**Figure 5.**
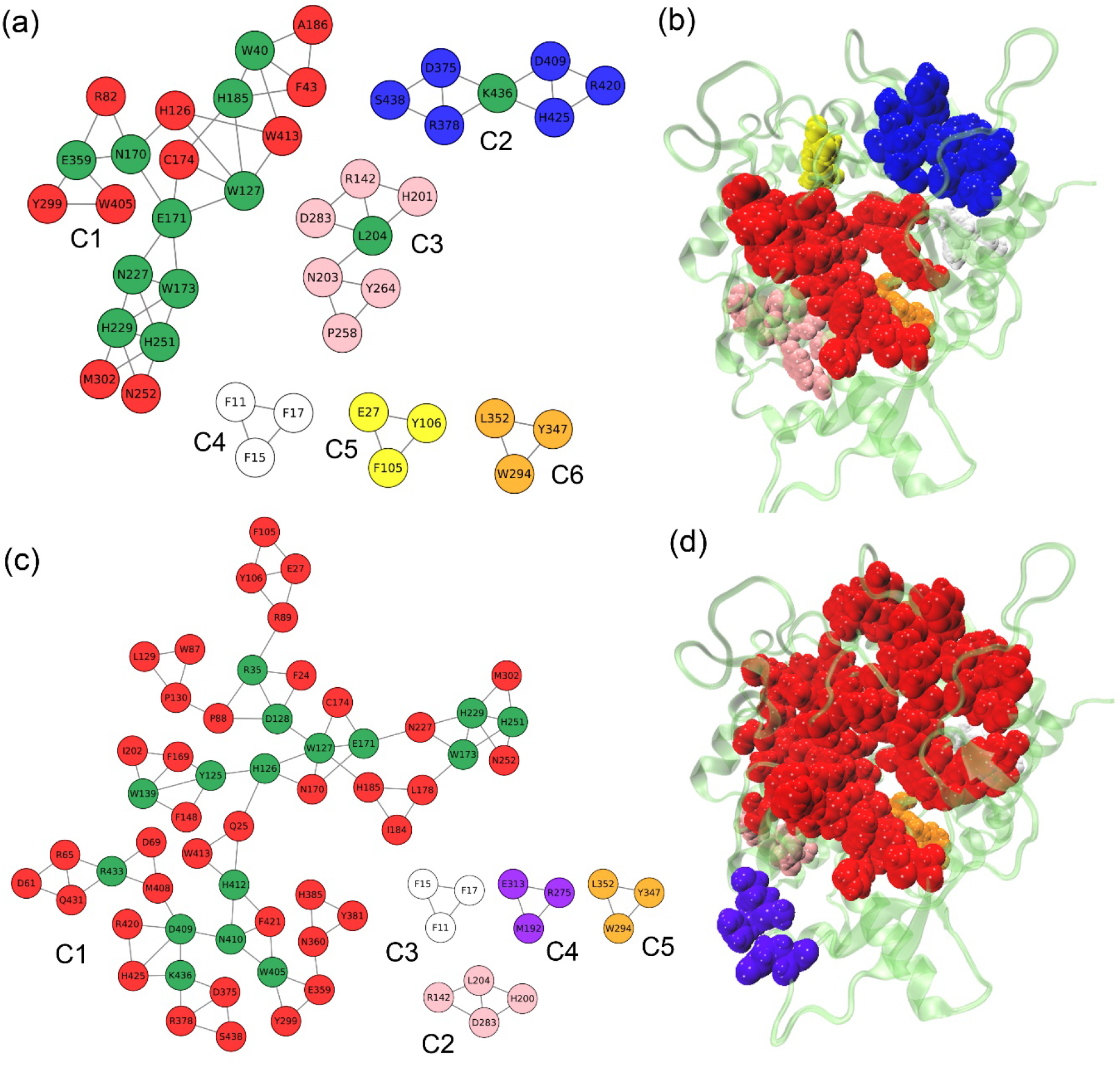
Variation of dynamically stable community structures of H0HC94 using protein structure network analysis. The network representation of the communities are shown in **(a)**, whereas **(b)** depicts the position of communities in the protein structure in S1 (WT+pNPGlc). The network representation of the communities in S3 (Q319A+pNPGlc) is presented in **(c)**, whereas **(d)** shows the position of the communities in the protein structure.

**Figure 6.**
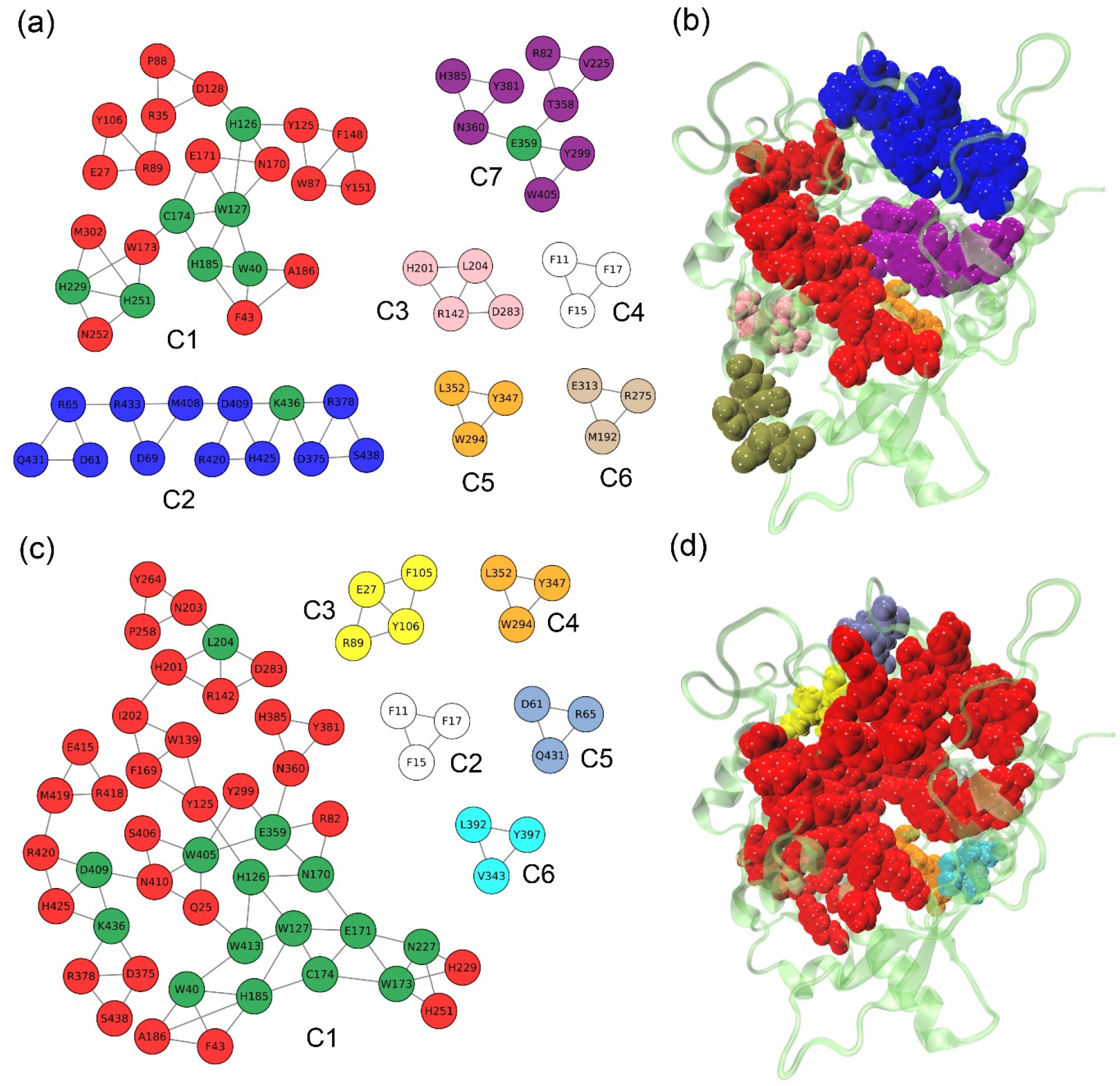
Variation of dynamically stable community structures of H0HC94 using protein structure network analysis. The network representation of the communities is shown in **(a)**, whereas **(b)** shows the position of the communities in the protein structure in S2 (WT+cellobiose). The network representation of the communities in S4 (Q319A+cellobiose) is presented in **(c)**, whereas **(d)** shows the position of the communities in the protein structure.

## Conclusion

Based on molecular docking results, we engineered Q319A H0HC94 by disrupting its interaction with glucose and enhancing cellobiose accessibility. We speculated that disrupting the enzyme-product (glucose) interaction at the entrance near the outside of the tunnel might facilitate the substrate (cellobiose or *p*NPGlc) access to the enzyme’s active site. The Q319A H0HC94 showed an enhancement of enzyme activity on cellobiose and increased cellobiose specificity. This study revealed that mutating a non-conserved residue (Q319) distant from the active site induced favorable conformational changes, leading to enhanced enzymatic activity. Our findings pave the way for engineering β-glucosidases without affecting the native active-site topology of the enzyme. This work also demonstrates the advantages of rational engineering of enzymes.

## Acknowledgement

SS is supported by a Junior Research Fellowship from IISER Kolkata, PC by an Inspire Fellowship, DST, Govt. of India, and BM by a IISER Kolkata Postdoctoral Fellowship.

## References

1. Goswami, S.; Nath, P.; Datta, S., Chapter 3 - Role of thermophilic cellulases and organisms in the conversion of biomass to biofuels. In Extremozymes and Their Industrial Applications, Arora, N. K.; Agnihotri, S.; Mishra, J., Eds. Academic Press: 2022; pp 85–113.

2. Aich, S.; Datta, S., Efficient Utilization of Lignocellulosic Biomass: Hydrolysis Methods for Biorefineries. In Biorefineries: A Step Towards Renewable and Clean Energy, Verma, P., Ed. Springer Singapore: Singapore, 2020; pp 273–295.

3. Datta, S., Recent Strategies to Overexpress and Engineer Cellulases for Biomass Degradation. Current Metabolomics 2016, 4 (1), 14–22.

4. Sinha, S. K.; Goswami, S.; Das, S.; Datta, S., Exploiting non-conserved residues to improve activity and stability of Halothermothrix orenii β-glucosidase. Appl Microbiol Biotechnol 2017, 101 (4), 1455–1463.

5. Lee, H. L.; Chang, C. K.; Jeng, W. Y.; Wang, A. H.; Liang, P. H., Mutations in the substrate entrance region of β-glucosidase from Trichoderma reesei improve enzyme activity and thermostability. Protein Eng Des Sel 2012, 25 (11), 733–40.

6. Santos, C. A.; Morais, M. A. B.; Terrett, O. M.; Lyczakowski, J. J.; Zanphorlin, L. M.; Ferreira-Filho, J. A.; Tonoli, C. C. C.; Murakami, M. T.; Dupree, P.; Souza, A. P., An engineered GH1 β-glucosidase displays enhanced glucose tolerance and increased sugar release from lignocellulosic materials. Sci Rep 2019, 9 (1), 4903.

7. Liu, X.; Cao, L.; Zeng, J.; Liu, Y.; Xie, W., Improving the cellobiose-hydrolysis activity and glucose-tolerance of a thermostable β-glucosidase through rational design. Int J Biol Macromol 2019, 136, 1052–1059.

8. Goswami, S.; Das, S.; Datta, S., Understanding the role of residues around the active site tunnel towards generating a glucose-tolerant β-glucosidase from Agrobacterium tumefaciens 5A. Protein Eng Des Sel 2017, 30 (7), 523–530.

9. Lenz, F.; Zurek, P.; Umlauf, M.; Tozakidis, I. E. P.; Jose, J., Tailor-made β-glucosidase with increased activity at lower temperature without loss of stability and glucose tolerance. Green chemistry 2020, 22 (7), 2234–2243.

10. Guo, B.; Amano, Y.; Nozaki, K., Improvements in Glucose Sensitivity and Stability of Trichoderma reesei β-Glucosidase Using Site-Directed Mutagenesis. PLoS One 2016, 11 (1), e0147301.

11. Cao, L. C.; Wang, Z. J.; Ren, G. H.; Kong, W.; Li, L.; Xie, W.; Liu, Y. H., Engineering a novel glucose-tolerant β-glucosidase as supplementation to enhance the hydrolysis of sugarcane bagasse at high glucose concentration. Biotechnol Biofuels 2015, 8, 202.

12. Cao, L.; Chen, R.; Huang, X.; Li, S.; Zhang, S.; Yang, X.; Qin, Z.; Kong, W.; Xie, W.; Liu, Y., Engineering of β-Glucosidase Bgl15 with Simultaneously Enhanced Glucose Tolerance and Thermostability To Improve Its Performance in High-Solid Cellulose Hydrolysis. J Agric Food Chem 2020, 68 (19), 5391–5401.

13. Goswami, S.; Gupta, N.; Datta, S., Using the β-glucosidase catalyzed reaction product glucose to improve the ionic liquid tolerance of β-glucosidases. Biotechnol Biofuels 2016, 9, 72.

14. Trott, O.; Olson, A. J., AutoDock Vina: improving the speed and accuracy of docking with a new scoring function, efficient optimization, and multithreading. J Comput Chem 2010, 31 (2), 455–61.

15. Wang, C.; Ye, F.; Chang, C.; Liu, X.; Wang, J.; Wang, J.; Yan, X. F.; Fu, Q.; Zhou, J.; Chen, S.; Gao, Y. G.; Zhang, L. H., Agrobacteria reprogram virulence gene expression by controlled release of host-conjugated signals. Proc Natl Acad Sci U S A 2019, 116 (44), 22331–22340.

16. Biovia, D. S., BIOVIA Discovery Studio. San Diego, 2020.

17. Schöning-Stierand, K.; Diedrich, K.; Fährrolfes, R.; Flachsenberg, F.; Meyder, A.; Nittinger, E.; Steinegger, R.; Rarey, M., ProteinsPlus: interactive analysis of protein–ligand binding interfaces. Nucleic Acids Research 2020, 48 (W1), W48–W53.

18. Brøns-Poulsen, J.; Petersen, N. E.; Hørder, M.; Kristiansen, K., An improved PCR-based method for site directed mutagenesis using megaprimers. Mol Cell Probes 1998, 12 (6), 345–8.

19. Case, D.; Belfon, K.; Ben-Shalom, I.; Brozell, S.; Cerutti, D.; Cheatham, T.; Cruzeiro, V.; Darden, T.; Duke, R.; Giambasu, G., AMBER 2020: University of California. San Francisco 2020.

20. Manna, B.; Ghosh, A., Molecular Insight into Glucose-Induced Conformational Change to Investigate Uncompetitive Inhibition of GH1 β-Glucosidase. ACS Sustainable Chemistry & Engineering 2021, 9 (4), 1613–1624.

21. Goswami, S.; Manna, B.; Chattopadhyay, K.; Ghosh, A.; Datta, S., Role of Conformational Change and Glucose Binding Sites in the Enhanced Glucose Tolerance of Agrobacterium tumefaciens 5A GH1 β-Glucosidase Mutants. J Phys Chem B 2021, 125 (33), 9402–9416.

22. Maier, J. A.; Martinez, C.; Kasavajhala, K.; Wickstrom, L.; Hauser, K. E.; Simmerling, C., ff14SB: Improving the Accuracy of Protein Side Chain and Backbone Parameters from ff99SB. J Chem Theory Comput 2015, 11 (8), 3696–713.

23. Kirschner, K. N.; Yongye, A. B.; Tschampel, S. M.; González-Outeiriño, J.; Daniels, C. R.; Foley, B. L.; Woods, R. J., GLYCAM06: a generalizable biomolecular force field. Carbohydrates. J Comput Chem 2008, 29 (4), 622–55.

24. Pettersen, E. F.; Goddard, T. D.; Huang, C. C.; Couch, G. S.; Greenblatt, D. M.; Meng, E. C.; Ferrin, T. E., UCSF Chimera--a visualization system for exploratory research and analysis. J Comput Chem 2004, 25 (13), 1605–12.

25. Martínez, L.; Andrade, R.; Birgin, E. G.; Martínez, J. M., PACKMOL: a package for building initial configurations for molecular dynamics simulations. J Comput Chem 2009, 30 (13), 2157–64.

26. Dolinsky, T. J.; Nielsen, J. E.; McCammon, J. A.; Baker, N. A., PDB2PQR: an automated pipeline for the setup of Poisson-Boltzmann electrostatics calculations. Nucleic Acids Res 2004, 32 (Web Server issue), W665–7.

27. Jorgensen, W. L.; Chandrasekhar, J.; Madura, J. D.; Impey, R. W.; Klein, M. L., Comparison of simple potential functions for simulating liquid water. The Journal of Chemical Physics 1983, 79 (2), 926–935.

28. Darden, T.; Perera, L.; Li, L.; Pedersen, L., New tricks for modelers from the crystallography toolkit: the particle mesh Ewald algorithm and its use in nucleic acid simulations. Structure 1999, 7 (3), R55–60.

29. Roe, D. R.; Cheatham, T. E., 3rd, PTRAJ and CPPTRAJ: Software for Processing and Analysis of Molecular Dynamics Trajectory Data. J Chem Theory Comput 2013, 9 (7), 3084–95.

30. Humphrey, W.; Dalke, A.; Schulten, K., VMD: visual molecular dynamics. J Mol Graph 1996, 14 (1), 33-8, 27-8.

31. Manna, B.; Ghosh, A., Understanding the conformational change and inhibition of hyperthermophilic GH10 xylanase in ionic liquid. Journal of Molecular Liquids 2021, 332, 115875.

32. Manna, B.; Ghosh, A., Structure and dynamics of ionic liquid tolerant hyperthermophilic endoglucanase Cel12A from Rhodothermus marinus. RSC Adv 2020, 10 (13), 7933–7947.

33. Felline, A.; Seeber, M.; Fanelli, F., PSNtools for standalone and web-based structure network analyses of conformational ensembles. Comput Struct Biotechnol J 2022, 20, 640–649.

34. Guo, B.; Sato, N.; Biely, P.; Amano, Y.; Nozaki, K., Comparison of catalytic properties of multiple β-glucosidases of Trichoderma reesei. Appl Microbiol Biotechnol 2016, 100 (11), 4959–68.

35. Xia, W.; Bai, Y.; Cui, Y.; Xu, X.; Qian, L.; Shi, P.; Zhang, W.; Luo, H.; Zhan, X.; Yao, B., Functional diversity of family 3 β-glucosidases from thermophilic cellulolytic fungus Humicola insolens Y1. Sci Rep 2016, 6, 27062.

36. Riou, C.; Salmon, J. M.; Vallier, M. J.; Günata, Z.; Barre, P., Purification, characterization, and substrate specificity of a novel highly glucose-tolerant beta-glucosidase from Aspergillus oryzae. Appl Environ Microbiol 1998, 64 (10), 3607–14.

37. Uchiyama, T.; Yaoi, K.; Miyazaki, K., Glucose-tolerant β-glucosidase retrieved from a Kusaya gravy metagenome. Front Microbiol 2015, 6, 548.

