## Supplementary Information for "Rational engineering of a β-glucosidase (H0HC94) from glycosyl family I (GH1) to improve catalytic performance on cellobiose"

**Figure S1. (a)** Cellobiose docked to H0HC94 (6RJO) by AutoDock vina.^1^ The structure was visualized by PyMOL.^2^ The interacting residues are shown in blue, and cellobiose is marked in green; **(b)** Interaction frequency distribution histogram of different H0HC94 residues with docked glucose molecule, calculated using in-house python script.

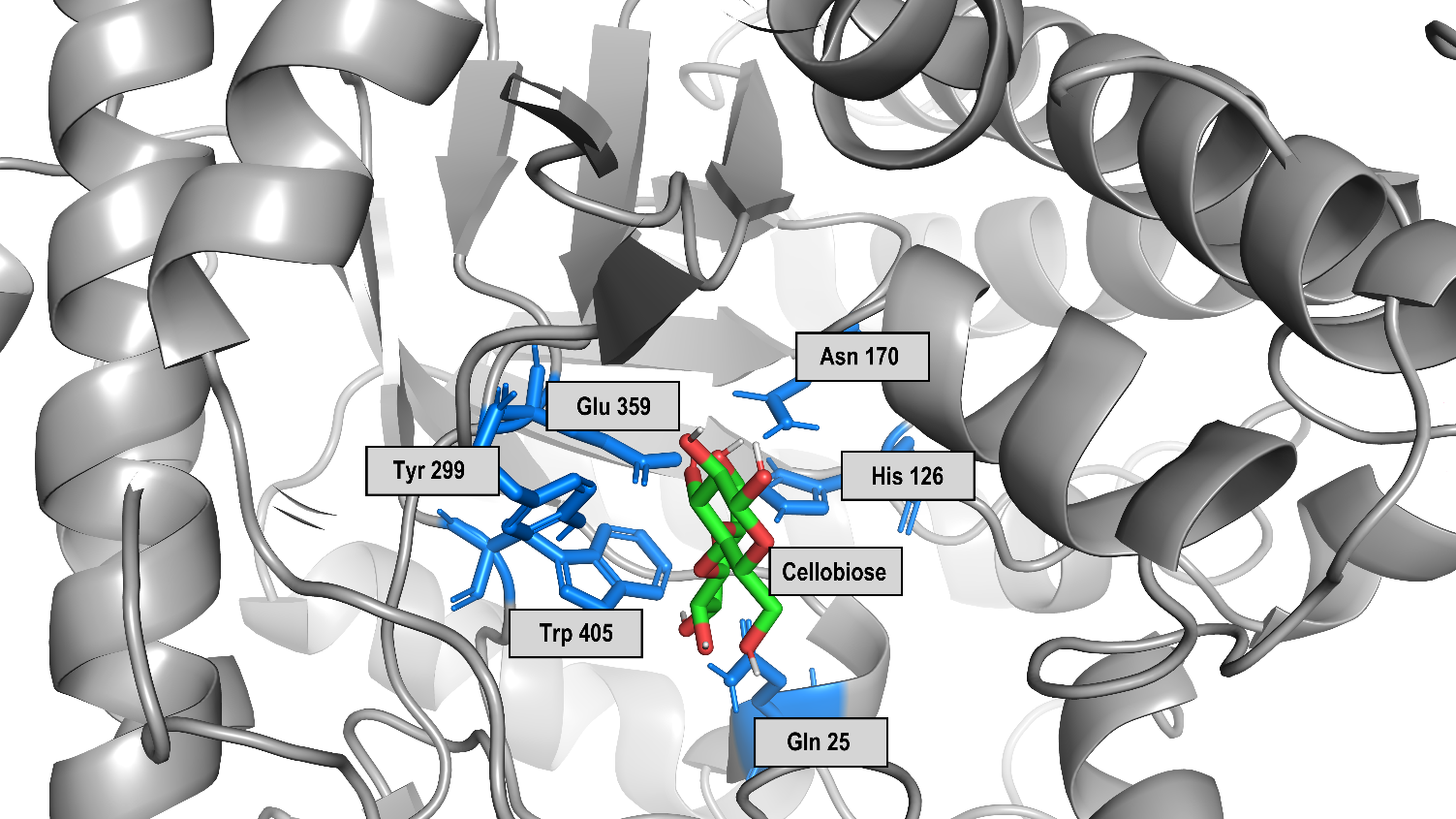
**(a)**

No. of interactions

| Amino acid residues in and around catalytic pocket of H0HC94 | No. of interactions Hydrogen bonds | No. of interactions Van der Waals interactions | No. of interactions Total no. of interactions |
| --- | --- | --- | --- |
| Q025 | 3 | 1 | 4 |
| H126 | 0 | 2 | 2 |
| W127 | 0 | 4 | 4 |
| N170 | 0 | 3 | 3 |
| E171 | 7 | 2 | 9 |
| W173 | 0 | 2 | 2 |
| C174 | 1 | 3 | 4 |
| L178 | 0 | 1 | 1 |
| I184 | 0 | 0 | 0 |
| H185 | 0 | 2 | 2 |
| N227 | 1 | 3 | 4 |
| A228 | 0 | 0 | 0 |
| H229 | 1 | 1 | 2 |
| S230 | 0 | 1 | 1 |
| N297 | 0 | 5 | 5 |
| Y298 | 0 | 3 | 3 |
| Y299 | 0 | 7 | 7 |
| T300 | 4 | 2 | 6 |
| P301 | 1 | 2 | 3 |
| M302 | 0 | 3 | 3 |
| R303 | 1 | 1 | 2 |
| Q319 | 2 | 1 | 3 |
| A322 | 0 | 2 | 2 |
| K327 | 2 | 0 | 2 |
| I330 | 0 | 0 | 0 |
| G331 | 1 | 1 | 2 |
| W332 | 0 | 7 | 7 |
| E333 | 1 | 0 | 1 |
| E359 | 1 | 2 | 3 |
| W405 | 2 | 1 | 3 |
| E412 | 1 | 1 | 2 |
| W413 | 1 | 1 | 2 |
| A414 | 0 | 0 | 0 |
| E415 | 0 | 0 | 0 |
| F421 | 0 | 2 | 2 |

**(b)**

**Figure S2.** 2D Interaction diagrams of glucose and active site tunnel residue Q319 of H0HC94. Made using PoseView tool.^3^

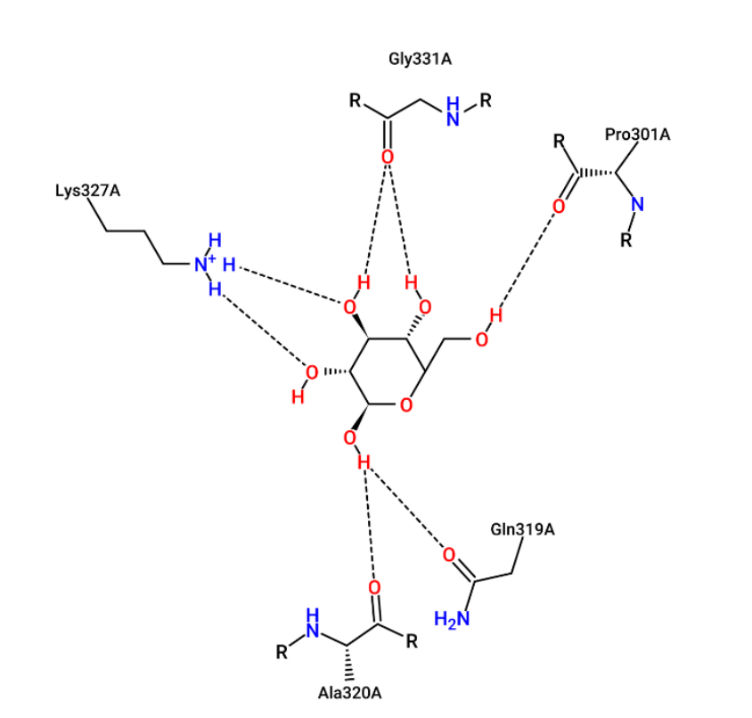

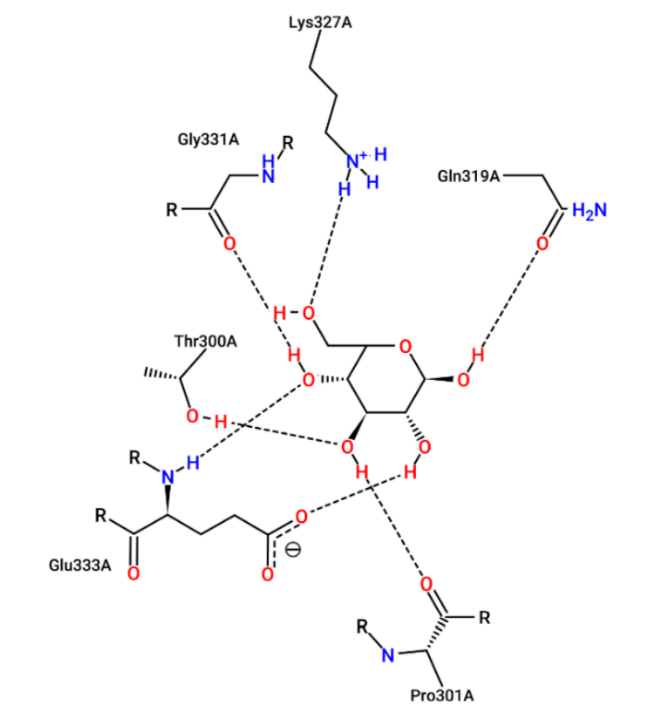

**(a)**

**(b)**

**Figure S3.** (a) The FPLC chromatogram of Q319A H0HC94 at 280 nm using 8 injection steps. (b) Desalting chromatogram of Q319A H0HC94 using two injection steps; (c) SDS-PAGE gel image of purified Q319A H0HC94, where the three lanes contain 12, 8 and 4 μg of purified protein (left to right) to check for impurities. The protein ladder (Thermo Scientific PageRuler Plus Prestained Protein Ladder, Thermo Fisher Scientific, Massachusetts, United States) was loaded in the last lane.

**(b)**

**(a)**

**(c)**

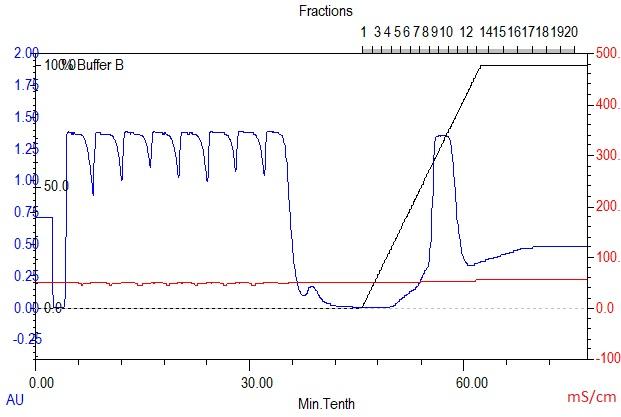

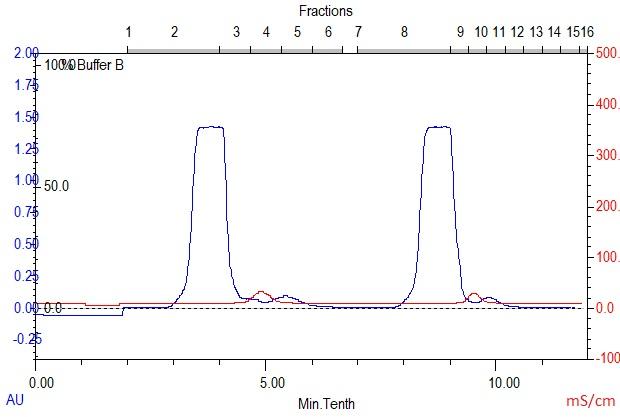

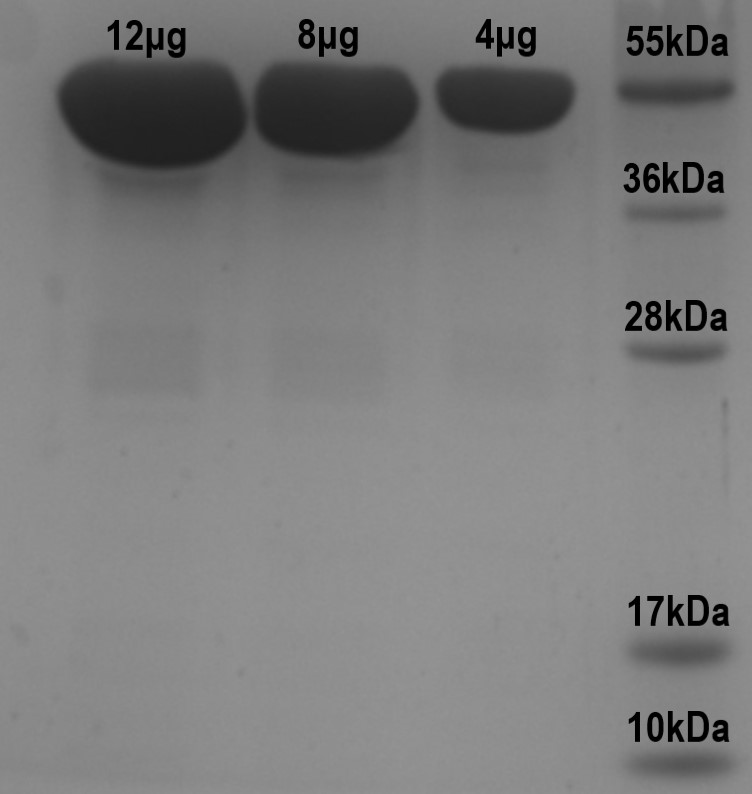

**Figure S4.** The ribbon representation of the H0HC94 structure. The active site catalytic residues (E171 and E359) and the Q319 are shown in the ball and stick model and coloured in red using Chimera.^4^

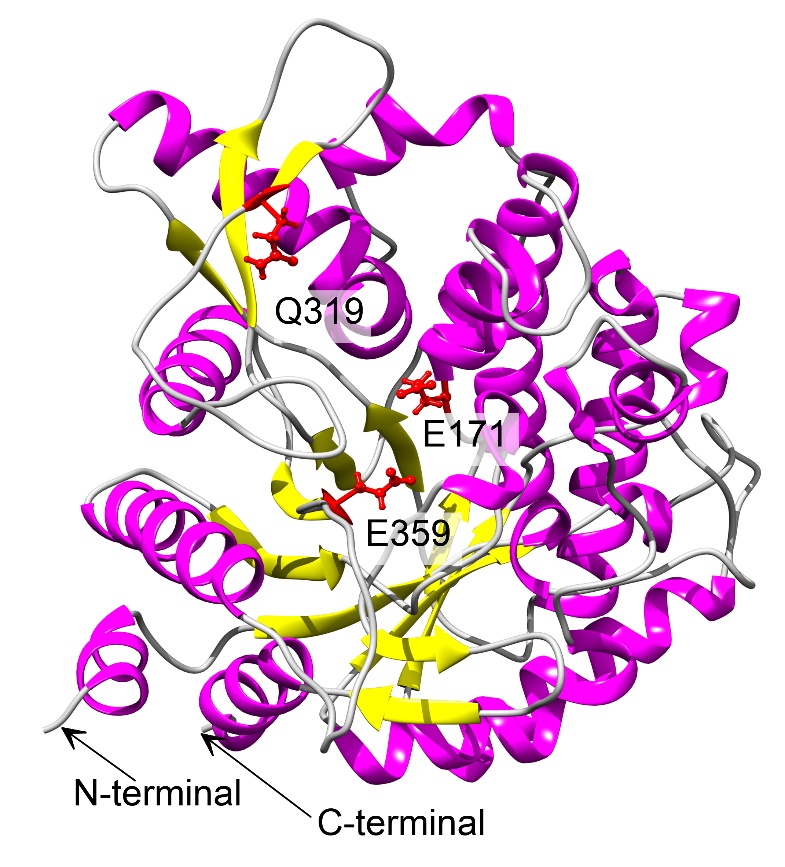

| **System Symbol** | **Composition** | **Water Molecules** | **Temperature (K)** | **pH** | **Simulation Time (ns)** |
| --- | --- | --- | --- | --- | --- |
| **S1** | WT + pNPGlc (0.02 M) | 14778 | 325 | 7.2 | 200 |
| **S2** | WT+ Cellobiose (0.02 M) | 14778 | 325 | 7.2 | 200 |
| **S3** | Q319A + pNPGlc (0.02 M) | 14778 | 322 | 7.0 | 200 |
| **S4** | Q319A + Cellobiose (0.02 M) | 14778 | 322 | 7.0 | 200 |

**Table S1.** System Details for MD simulation

**Table S2.** Secondary structure analysis by fitting the circular dichroism (CD) data of Q319A H0HC94 and WT. The secondary structure percentages were determined using the Beta Structure Selection (BeStSel) webserver.^5^

| **Secondary Structure (%)** | **H0HC94** | **Q319A H0HC94** |
| --- | --- | --- |
| **Alpha helix** | 17.4 | 14.8 |
| **Antiparallel β-sheets** | 31.45 | 25.6 |
| **Parallel β-sheets** | 0.0 | 0.0 |
| **Turns** | 13.0 | 11.1 |
| **Others** | 38.3 | 48.5 |

**Table S3.** Differential fluctuation of active site tunnel residues

| **Position** | **Residue** | **S1**  **(Å)** | **S2**  **(Å)** | **S3**  **(Å)** | **S4**  **(Å)** |
| --- | --- | --- | --- | --- | --- |
| **Active site**  **Residues** | **E171**  **E359** | 0.6783  0.4154 | 0.5211  0.4323 | 0.5100  0.4546 | 0.8048  0.4459 |
| **Tunnel**  **Residues** | **H126**  **W127**  **N170**  **C174**  **N297**  **W331**  **W405**  **W413** | 0.4689  0.5198  0.6178  0.5111  0.4462  0.7884  0.4206  0.9444 | 0.5337  0.5285  0.5777  0.4943  0.4651  0.8472  0.4281  0.9155 | 0.3846  0.3898  0.4391  0.4562  0.5407  0.7645  0.4054  0.8808 | 0.5279  0.6021  0.6347  0.5025  0.4648  0.7618  0.4078  1.0734 |
| **Gatekeeper**  **Residues** | **L178**  **H185**  **N227**  **H229**  **Y299**  **T300** | 0.6192  0.6785  0.4708  0.5274  0.4869  0.5397 | 0.4900  0.5970  0.8504  0.5249  0.5119  0.5178 | 0.4767  0.4894  0.8422  0.6200  0.5567  0.5688 | 0.6267  0.6032  0.5857  0.4602  0.5223  0.5631 |

**References**

1.	Trott, O.; Olson, A. J., AutoDock Vina: improving the speed and accuracy of docking with a new scoring function, efficient optimization, and multithreading. *J Comput Chem* **2010,** *31* (2), 455-61.

2.	Schrödinger, L., The PyMOL Molecular Graphics System. 2022.

3.	Schöning-Stierand, K.; Diedrich, K.; Fährrolfes, R.; Flachsenberg, F.; Meyder, A.; Nittinger, E.; Steinegger, R.; Rarey, M., ProteinsPlus: interactive analysis of protein–ligand binding interfaces. *Nucleic Acids Research* **2020,** *48* (W1), W48-W53.

4.	Pettersen, E. F.; Goddard, T. D.; Huang, C. C.; Couch, G. S.; Greenblatt, D. M.; Meng, E. C.; Ferrin, T. E., UCSF Chimera--a visualization system for exploratory research and analysis. *J Comput Chem* **2004,** *25* (13), 1605-12.

5.	Micsonai, A.; Moussong, É.; Wien, F.; Boros, E.; Vadászi, H.; Murvai, N.; Lee, Y. H.; Molnár, T.; Réfrégiers, M.; Goto, Y.; Tantos, Á.; Kardos, J., BeStSel: webserver for secondary structure and fold prediction for protein CD spectroscopy. *Nucleic Acids Res* **2022,** *50* (W1), W90-8.
